# CD6 Regulates CD4 T Follicular Helper Cell Differentiation and Humoral Immunity During Murine Coronavirus Infection

**DOI:** 10.1101/2024.07.26.605237

**Authors:** Amber Cardani-Boulton, Feng Lin, Cornelia C Bergmann

## Abstract

During activation the T cell transmembrane receptor CD6 becomes incorporated into the T cell immunological synapse where it can exert both co-stimulatory and co-inhibitory functions. Given the ability of CD6 to carry out opposing functions, this study sought to determine how CD6 regulates early T cell activation in response to viral infection. Infection of CD6 deficient mice with a neurotropic murine coronavirus resulted in greater activation and expansion of CD4 T cells in the draining lymph nodes. Further analysis demonstrated that there was also preferential differentiation of CD4 T cells into T follicular helper cells, resulting in accelerated germinal center responses and emergence of high affinity virus specific antibodies. Given that CD6 conversely supports CD4 T cell activation in many autoimmune models, we probed potential mechanisms of CD6 mediated suppression of CD4 T cell activation during viral infection. Analysis of CD6 binding proteins revealed that infection induced upregulation of *Ubash3a,* a negative regulator of T cell receptor signaling, was hindered in CD6 deficient lymph nodes. Consistent with greater T cell activation and reduced UBASH3a activity, the T cell receptor signal strength was intensified in CD6 deficient CD4 T cells. These results reveal a novel immunoregulatory role for CD6 in limiting CD4 T cell activation and deterring CD4 T follicular helper cell differentiation, thereby attenuating antiviral humoral immunity.

**Importance:** CD6 monoclonal blocking antibodies are being therapeutically administered to inhibit T cell activation in autoimmune disorders. However, the multifaceted nature of CD6 allows for multiple and even opposing functions under different circumstances of T cell activation. We therefore sought to characterize how CD6 regulates T cell activation in the context viral infections using an *in vivo* murine coronavirus model. In contrast to its role in autoimmunity, but consistent with its function in the presence of superantigens, we found that CD6 deficiency enhances CD4 T cell activation and CD4 T cell help to germinal center dependent antiviral humoral responses. Finally, we provide evidence that CD6 regulates transcription of its intracellular binding partner UBASH3a, which suppresses T cell receptor signaling and consequently T cell activation. These findings highlight the context dependent flexibility of CD6 in regulating *in vivo* adaptive immune responses, which may be targeted to enhance anti-viral immunity.

## Introduction

T cell activation through engagement of the T cell receptor (TCR) is critical to combat pathogens and tumors but can also cause detrimental injury if left unchecked. For this reason, T cell activation is a highly complex process involving multiple layers of regulation that are dependent on the concise orchestration of numerous signaling and scaffolding proteins. Consequently, the diverse composition of the TCR signalosome allows for significant flexibility in the strength of the TCR signal, which, rather than a simple binary “yes or no” signal, the strength of the TCR governs T cell activation, survival, and functional differentiation^1–4^.

The intensity of the TCR signal is fine-tuned by, among other mechanisms, TCR co-receptors that can enhance (co-stimulatory) or dampen (co-inhibitory) the TCR signal strength. CD6 is a cell-surface glycoprotein that has been demonstrated to function as both a TCR co-stimulatory or a co-inhibitory receptor during T cell activation in a context dependent manner^5,6^. The functional heterogeneity of CD6 resides in its cytoplasmic tail, which constitutes one of the longest leukocyte intracellular domains, thus making it challenging to characterize the CD6 signaling cascade and identify mechanisms by which it regulates TCR signaling ^5–7^. A number of CD6 intracellular binding proteins that positively propagate the TCR signal, most notably SLP76 and Zap70, have been confirmed. Conversely, few CD6 signaling proteins that negatively regulate TCR signaling have been identified. However, Ubiquitin associated and SH3 domain containing A (UBASH3a), formally known as suppressor of T cell signaling 2 (STS-2), was recently determined to directly interact with the cytoplasmic tail of CD6 ^5–7^.

While characterization of the CD6 signaling pathways remains incomplete, CD6 polymorphisms have been linked to either the susceptibility or severity of multiple autoimmune diseases^9–11^. Furthermore, in preclinical murine models of autoimmunity, inhibition of CD6 signaling was shown to prevent TCR co-stimulation, thereby limiting activation of autoreactive T cells^15^. Therefore, monoclonal CD6 blocking antibodies have been developed as clinical therapeutic treatments^8^. To this point, the anti-CD6 monoclonal antibody Itolizumab has been approved and is in use clinically in India for the treatment of psoriasis^8^. CD6 blockade has also shown promise in clinical trials for the treatment of rheumatoid arthritis and has recently gained interest as a potential cancer therapy^12–14^. However, despite indications that CD6 has a suppressive role in the presence of bacterial superantigens, the function of CD6 in the context of infectious disease remains under-studied^16^.

Given the opposing functions of CD6 in different disease and clinical settings, we sought to determine how CD6 regulates T cell activation in an established model of viral encephalomyelitis. The attenuated recombinant neurotropic murine beta coronavirus MHV-A59 (mCoV), was chosen for this study as both the CD4 T helper 1 (TH1) cells and cytotoxic CD8 T cells are essential for the control of infectious virus within the CNS^17–20^. Furthermore, the generation of mCoV-specific antibodies is dependent on the CD4 follicular helper (T_FH_) cells, which mediate both germinal center (GC) formation as well as somatic-hypermutation of the B cell immunoglobulin variable chain, thus enabling the generation of high-affinity class switched antigen-specific B cells^20–22^.

The study herein revealed that the absence of CD6 resulted in greater CD4 T cell activation in the CNS draining cervical lymph nodes (cLNs) following mCoV infection. An increase in total CD4 T cell numbers was accompanied by more pronounced differentiation into CD4 T_FH_ cells. As a result, cLN GC reactions were accelerated, complemented by the rapid appearance of high-affinity mCoV-specific antibodies in the serum of CD6 knockout (KO) mice. Increased CD4 T cell activation in the absence of CD6 was associated with impaired transcriptional upregulation of the established negative regulator of TCR signaling *Ubash3a* ^6–7^. In agreement with increased CD4 T cell activation and decreased UBASH3a activity, the intensity of the TCR signal was greater in CD6 deficient CD4 T cells. These data are the first to link CD6 with transcriptional upregulation of *Ubash3a* and reveal pivotal novel roles of CD6 as a negative regulator of antiviral CD4 T cell activation, CD4 T_FH_ cell differentiation, and antiviral humoral responses. Overall, these results highlight the context dependent functions of CD6 in regulating adaptive immune responses.

## Materials and Methods

### Mice and Infections

All procedures involving mice were approved by the Institutional Animal Care and Use Committee of Cleveland Clinic and carried out in accordance with the US Department of Health and Human Services Guide for the Care and Use of Laboratory Animals and institutional guidelines. CD6 KO mice on the DBA-1 background were generated in the laboratory of Dr. Feng Lin and maintained under pathogen-free conditions in the Cleveland Clinic Lerner Research Institute animal facility ^15^. WT controls were also maintained onsite and housed in the same room with occasional supplementation as well as rejuvenation of the breeders with mice purchased from Jackson Laboratory (strain number 000670). As previously reported^17, 21–23^ 6–8-week-old gender matched male and female WT and CD6 KO mice were intracranially (IC) infected with 10,000 PFU of the recombinant mCoV strain MHV-A59 whose non-essential open reading frame of gene 4 had been replaced with enhanced green fluorescent protein and was generously provided by Dr. Das Sarma^24^.

### Flow cytometry

The CNS draining deep cLN, brains, and spinal cords were isolated from phosphate buffer saline perfused mice. Tissues were finely minced, and single-cell suspensions obtained after mechanical homogenization through a 70micron strainer. Myelin was removed from CNS tissue by centrifugation at 850g for 45min at 4°C in 30% Percoll (Cytiva 17089101). After washing in 1X PBS, cells were resuspended in FACS buffer (1X PBS, 1% BSA, +/-0.1% NaN3) and stained with fluorescently conjugated antibodies in the presence of FC block (clone 2.4G2) for 30min at 4 degrees. The following antibodies were used: CD45 (clone 30-F11), CD3 (clone 17A2), CD4 (clone RM4-5), CD8 (clone 53-6.7), CD44 (clone IM7), CXCR5 (clone 2G8), PD1 (clone 29F.1A12), CD6 (J90-462), CD19 clone (clone 1D3), IgD (clone 11-26c.2a), GL7 (clone GL7), and CD138 (clone 281-2). After staining, cells were washed and either resuspended in FACS buffer containing DAPI dye for immediate analysis or stained with fixable live/dead dyes (Invitrogen Catalog number: L34957 or Beckman Coulter Catalog number: C36628) according to manufacturer’s protocol followed by fixation in 4% PFA. The eBioscience FoxP3/ Transcription Factor Staining Kit was used for intracellular staining according to the manufactures protocol. The following antibodies were used: BCL6 (clone K112-91), Tbet (clone ebio4B10), and IRF4 (Clone IRF.3E4) in the presence of FC block. Cells were collected using a 6-laser Beckman CytoFLEX LX. The resulting data were compensated and analyzed with FlowJo software (Tree Star, Inc., Ashland, OR) using the gating strategy exemplified in Supplemental Figure 1.

### Immunofluorescence

The CNS draining deep cLN were isolated from phosphate buffer saline perfused mice and snap-frozen in Tissue-Tek O.C.T (Fisher). 10micron slices were obtained using a Leica CM3050 cryostat and slide-mounted. Sections were fixed in 4% PFA and permeabilized with Triton X-100. After blocking, cLNs were stained with anti-CD3 (clone 17A2), GL7 (clone GL7), and B220 (clone RA3-6B2).

Corresponding secondary antibodies were used as necessary and sections were mounted using ProLong Gold Antifade Mountant with DNA Stain DAPI. Entire cLNs were scanned at 20X or 40X magnification using a Leica DM6B upright microscope equipped for Fluorescence and Brightfield microscopy. Images were analyzed using Image J software (NIH; http://rsbweb.nih.gov/ij) implementing the FIJI plugin set (http://pacific.mpi-cbg.de/wiki/index.php/Fiji).

### Quantitative real-time PCR

The cLNs, brains, and spinal cords were isolated from phosphate buffer saline perfused mice and immediately placed in trizol or Qiagen RLT buffer on ice. Tissue was homogenized using the Qiagen TissueLyser with stainless-steel beads and stored at -80°C. RNA was isolated according to the manufacturer’s instructions. Samples were DNaseI treated (Invitrogen Catalog number: 18068015) according to the manufacturer’s instructions and cDNA was synthesized using MMLV reverse transcriptase (Invitrogen Catalog number: 28025021). Quantitative real-time PCR was performed using PowerUp SYBR Green Master Mix (Fisher A25742) on a 7500 fast real-time PCR system (Applied Biosystems, Foster City, CA). Transcript levels were calculated relative to the levels of the *Gapdh* housekeeping gene using the following cycle threshold (C T) formula: 2 ^[CT(*Gapdh*) - CT(target gene)].

Primers for glyceraldehyde 3-phosphate dehydrogenase (*Gapdh*), activation-induced cytidine deaminase (*Aicda*), T-box transcription factor 21 (*Tbx21*), and ubiquitin associated and SH3 domain containing A (*UBASH3a*) were purchased for Sybr Green analysis from Qiagen (Catalog number: 330001). All other primers are as follows: *Il17* (F: CTCCACCGCAATGAAGAC and R: CTTTCCCTCCGCATTGAC), *Foxp3* (F: CTGCTCCTCCTATTCCCGTAAC and R: AGCTAGAGGCTTTGCCTTCG), *Ifng* (F: CCAAGTTTGAGGTCAACAACCC and R: AACAGCTGGTGGACCACTC), mCoV-*N* (F: GCCAAATAATCGCGCTAGAA and R: CCGAGCTTAGCCAAAACAAG), *Irf4* F: (GAACGAGGAGAAGAGCGTCTTC and R: GTAGGAGGATCTGGCTTGTCGA)

### mCoV-specific IgG ELISA and affinity index50

As previously described^25–26^, serum collected from individual infected mice was serially diluted across virus-coated ELISA plates. After incubation and washing, mCoV-specific antibodies bound to virus coated plates were detected using HRP conjugated anti-mouse IgG antibodies, TMB substrate, and ELISA stop solution. A dilution in which the fluorescence intensity was well within the linear curve was used to determine differences between WT and CD6 KO mice using optical density. Naïve controls were used to determine background signal. For tissue antibody detection, whole tissue was placed in ice cold PBS and homogenized using a Dounce Homogenizer. Cells and debris were removed by centrifugation and the supernatant stored at -80° C prior to serial dilution as performed with serum samples. Data was plotted across all dilutions tested.

At a concentration that was experimentally determined using the above mCoV-specific ELISA, samples were incubated on virus-coated plates overnight. After washing, serial dilutions of ammonium thiocyanate (3M-0M) were added to each sample and incubated for 15min on a shaker at room temperature. After washing mCoV-specific antibodies still bound to the virus coated plate were detected using HRP conjugated anti-mouse IgG antibodies, TMB substrate, and ELISA stop solution. The Affinity index 50 was determined as the concentration of ammonium thiocyanate at which 50% of the antibody signal was lost^27–29^.

### Statistics

GraphPad Prism software was used to plot data points with SEM for error bars. Statistical significance was determined using GraphPad Prism Software as specified in the figure legends.

## Results

### CD6 suppresses adaptive immune cell expansion and activation after mCoV infection

To assess a potential role for CD6 in regulating T cell activation during viral encephalomyelitis CD6 KO and wild type (WT) control mice were intracranially (IC) infected with an attenuated recombinant murine Coronavirus (mCoV) strain MHV-A59^24^. In the absence of CD6 there was greater expansion of the CD4 T cells in the cLNs as early as day 4 post infection (PI), which was sustained at day 7 PI (Figure 1A). In addition, a greater percentage of CD4 T cells expressed high levels of the activation marker CD44 (Figure 1B). Examination of the CD8 T cell population revealed transiently increased expansion at day 4PI that contracted to WT levels by day 7 PI (Figure 1C), although increased activation was evident at both days 4 and 7PI (Figure 1D). Importantly, no significant differences in T cell numbers or activation statuses were observed in cLNs of naïve CD6 KO compared to WT mice (Figure 1A-D). Interestingly, the total number of B cells in the CD6 KO cLNs was also elevated at day 7PI (Figure 1E). However, within the cLNs and brain CD6 was only detectable on CD4 and CD8 T cells despite reports that CD6 is expressed on B1a B cells and some CD56 expressing NK cells^30, 31^ (Supplemental Figure 2A-D). Importantly, the increased lymphocyte activation in the CD6 KO cLNs could not be attributed to increased viral loads as transcripts of the mCoV nucleocapsid protein (N) were similar between WT and CD6 KO cLNs (Figure 1F). Therefore, inhibition of CD6 signaling resulted in greater CD4 T cell activation and expansion, followed by increased B cell accumulation in the cLNs after CNS infection with mCoV. These data contrasted studies of CD6 function in experimental autoimmune models, but are consistent with *in vitro* analysis of CD6 during bacterial superantigen exposure^15–16, 46–47^.

**Figure 1:**
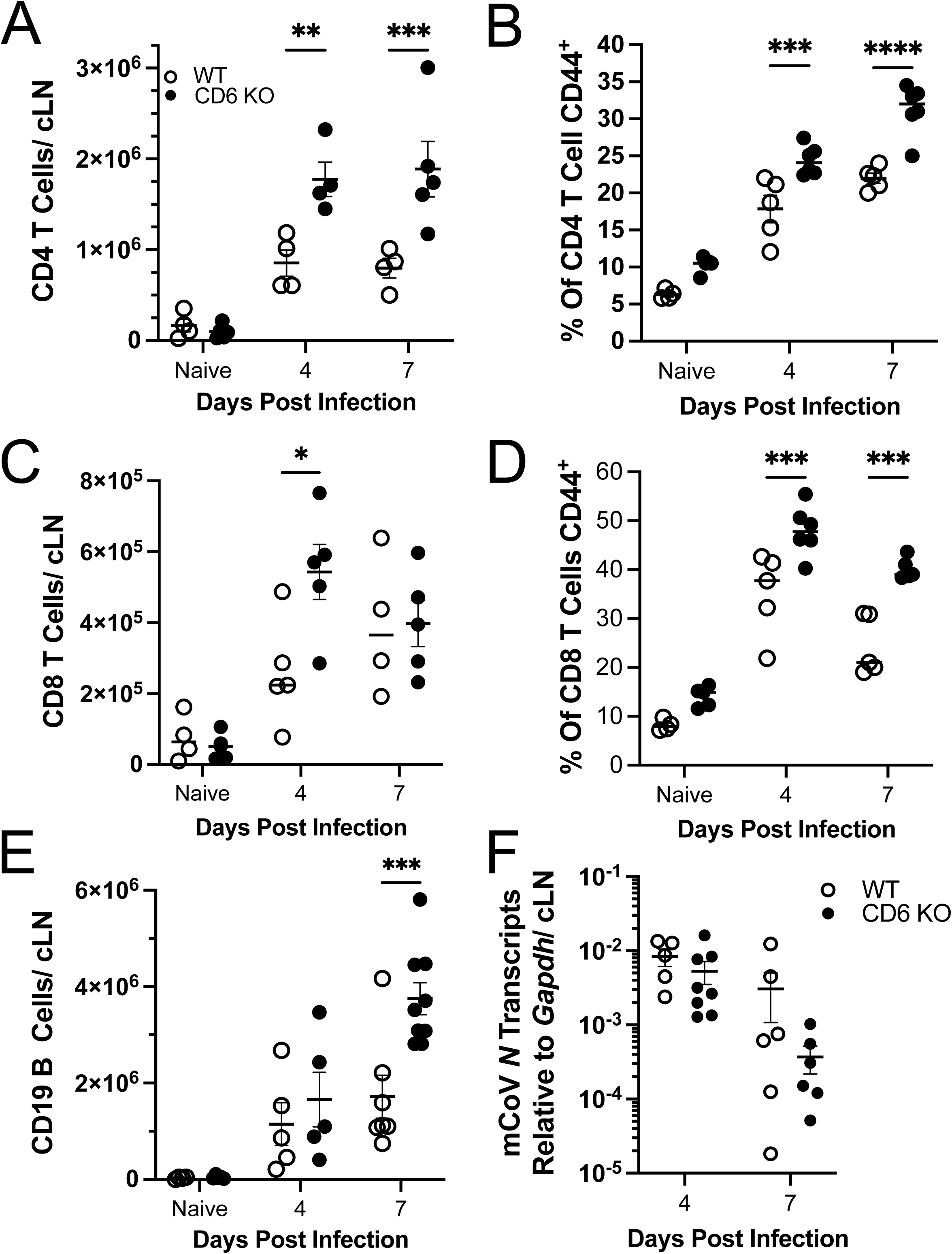
CD6 deficiency results in enhanced adaptive immune cell expansion and activation in the cervical lymph nodes after mCoV infection. cLNs from WT and CD6 KO mice infected with 10,000 PFU of mCoV were taken at the indicated time points and analyzed by flow cytometry for the A) total number of CD4 T cells, B) percent of CD4 T cells that are CD44 high, C) total number of CD8 T cells, D) percent of CD8 T cells that are CD44 high, and E) total number of CD19 B cells. cLNs were also analyzed by F) qRT-PCR for transcripts of the mCoV nucleocapsid gene (N). Each data point represents a single mouse and a minimum of two individual experiments were pooled for each time point. Significance was determined using a Two-way ANOVA with a Bonferroni’s post-hoc test and denoted as * for p<0.05, ** for p<0.01, *** for p<0.001,and **** for p<0.0001.

### CD6 regulates CD4 T helper cell differentiation

Given the expansion of cLN B cells, which do not express CD6, we next examined if CD4 T_FH_ differentiation was altered in CD6 KO cLNs. Flow cytometry analysis revealed that, even when normalized for the increase in total CD4 T cell numbers, there was increased differentiation of CD4 T cells into T_FH_ cells (PD1^+^, CXCR5^+^) as early as day 4PI, which was sustained through at least day 7PI in CD6 KO cLNs (Figure 2A). CD4 T cells expressing BCL6, the transcription factor essential for T_FH_ cell differentiation, was also elevated in CD4 T cells from CD6 KO cLNs compared to WT cLNs at day 4 and 7PI (Figure 2B)^32^. Conversely, the percent of CD4 T cells expressing T-bet, the transcription factor essential for TH1 differentiation, was similar between WT and CD6 KO mice^33^ (Figure 2C). Therefore, the increase in total T-bet^+^ CD4 T cells was consistent with the overall increase in total CD4 T cells and not greater skewing towards TH1 differentiation. There was also no significant difference in *Ifng* transcript levels indicating that the CD4 TH1 effector response was not significantly altered in CD6 KO mice (Figure 2D). *Il17* mRNA could not be detected in either WT or CD6 KO cLNs (data not shown), and no difference in *Foxp3* mRNA transcripts was observed (Figure 2E). Taken together these data demonstrate that CD6 is able to limit CD4 T_FH_ cell differentiation in addition to suppressing overall T cell activation and expansion following mCoV infection.

**Figure 2:**
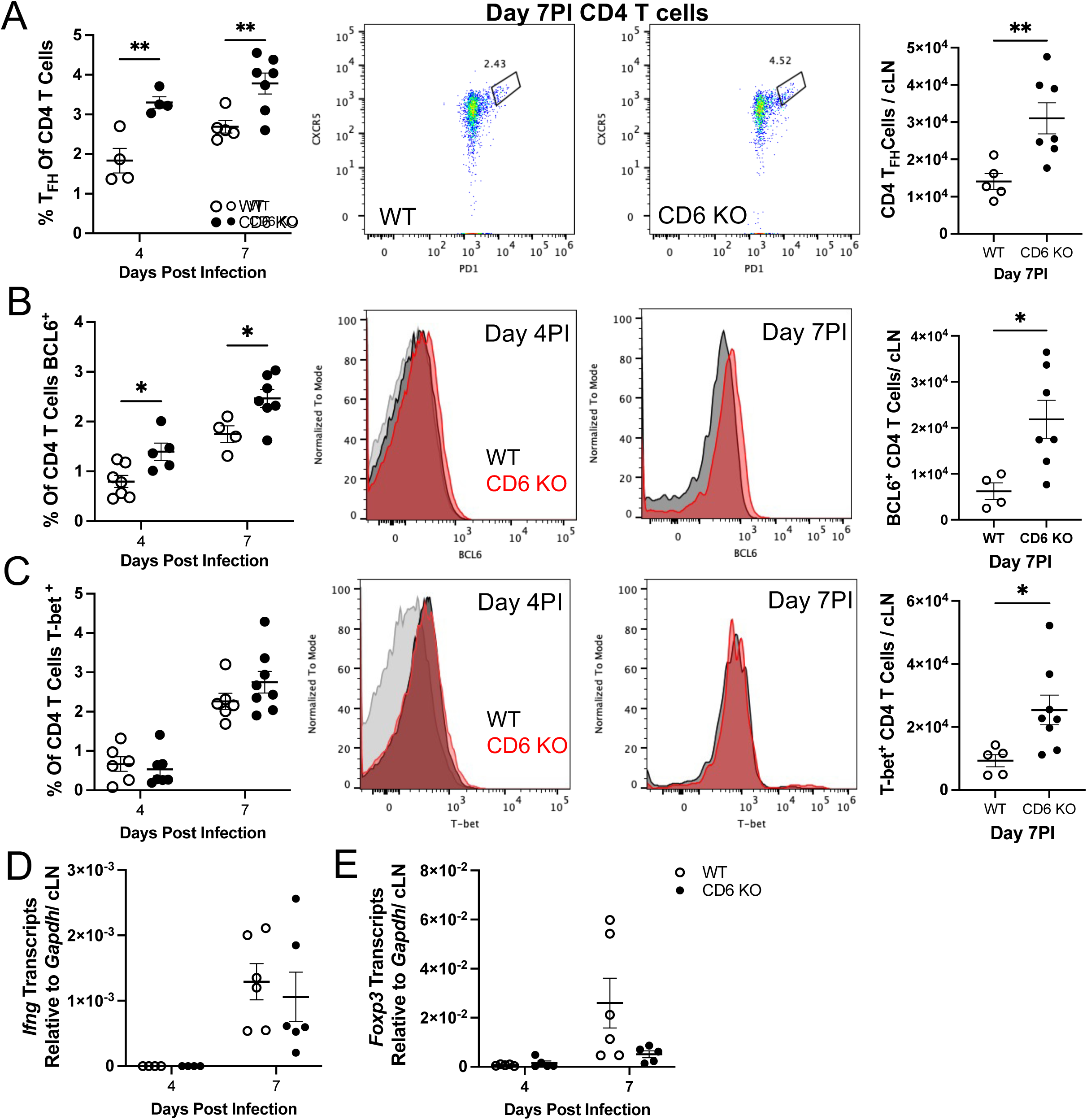
CD6 regulates CD4 T helper cell differentiation in cervical lymph nodes after mCoV infection. cLNs from mCoV infected WT and CD6 KO mice were taken at the indicated time points and analyzed by flow cytometry for the proportion and number of CD4 T cells that are **A)** T_FH_ cells (CXCR5^+^, PD1^+^), **B)** BCL6^+^, and **C)** T-Bet^+^. cLNs s were also analyzed by qRT-PCR for **D)** *Ifng* and **E)** *Foxp3* transcripts. Each data point represents a single mouse with experiments pooled from a minimum of two independent experiments for each time point. Flow cytometry plots and histograms are of a representative WT(black) and CD6 KO (red) mouse. FMO staining controls (gray) were included in the day 4PI histograms. Significance was determined using a Two-way ANOVA with a Bonferroni’s post-hoc test or an unpaired T-test and denoted as * for p<0.05, ** for p<0.01, *** for p<0.001,and **** for p<0.0001.

### GC differentiation is enhanced in CD6 KOs during infection

To our knowledge this is the first time that CD6 has been linked to regulating T_FH_ cell differentiation. We therefore analyzed the *in vivo* effector capacity of CD4 T_FH_ cells generated in the CD6 KO mice by examining cLN GC formation. Subsequent to the appearance of CD4 T_FH_ cells in the CD6 KO cLNs, a larger fraction of B cells in the CD6 KO cLNs had downregulated cell surface IgD, indicative of early activation (Figure 3A). Similarly, a greater proportion of the already enlarged CD19 B cell population also expressed the GC B cell marker GL7 at day 7 PI in CD6 KO cLNs (Figure 3B).

**Figure 3:**
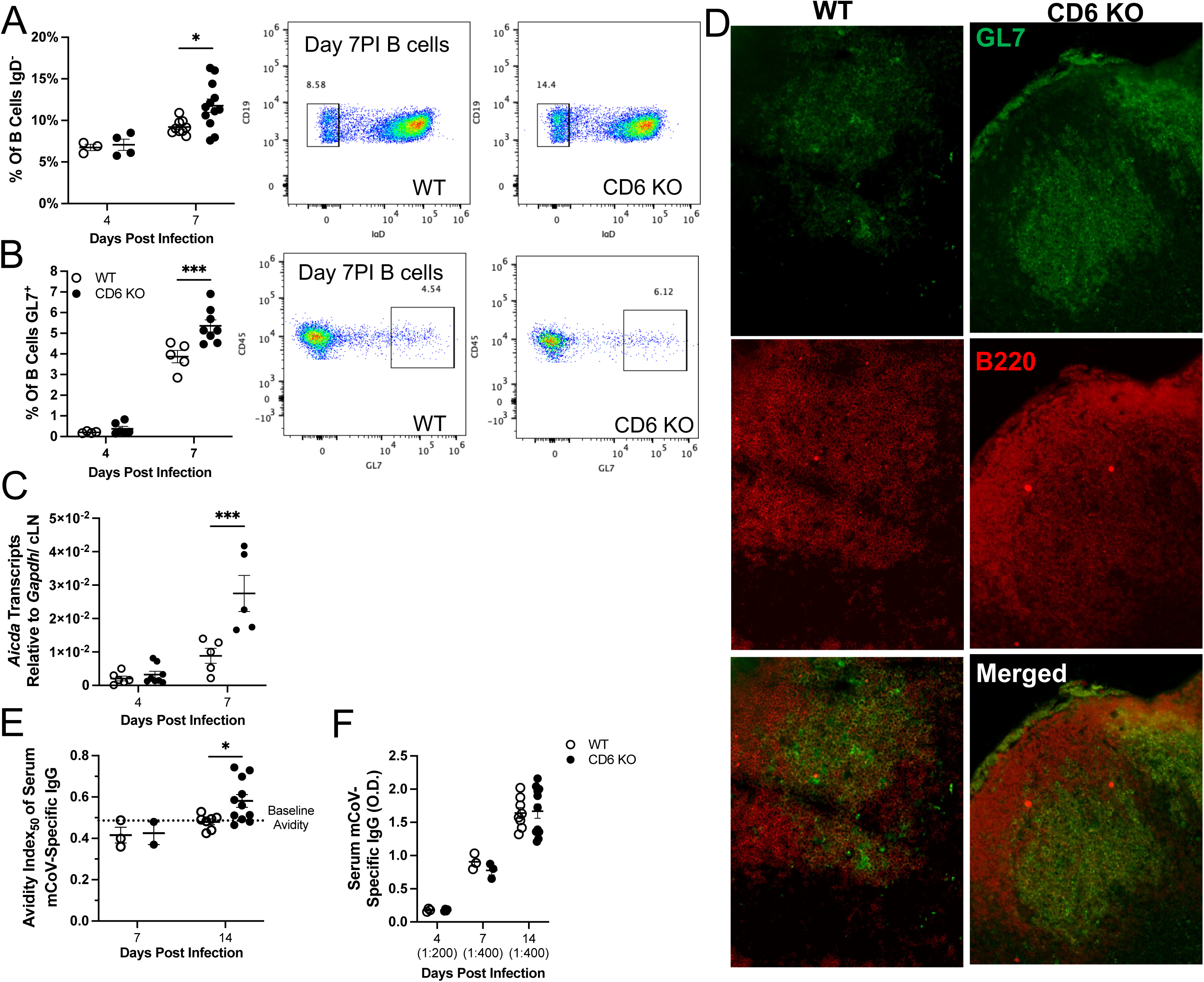
Germinal centers are enhanced in the absence of CD6. WT and CD6 KO mice were infected with mCoV. At day 4 and 7PI cLN CD19^+^ B cells were examined by flow cytometry for expression of **A)** IgD or **B)** GL7 (denoting germinal center B cells). Additionally, qRT-PCR was used to analyze expression of **C)** *Aicda* in cLNs. Day 14PI flash fozen cLNs were **D**) stained for B220 (red), GL7 (green), and CD3 (blue) to confirm GC formation within B cell zones. Serum collected at days 4, 7, and 14 PI was analyzed for mCoV-specific IgG **E)** avidity index_50_ and **F)** titers by ELISA. “()” indicates the experimentally determined concentration of the serum dilution used for the analysis. Each data point represents a single mouse and a minimum of two experiments were pooled for each time point. Flow plots are of a representative mouse. Significance was determined by Two-way ANOVA using a Bonferroni’s post-hoc test and denoted as * for p<0.05, ** for p<0.01, *** for p<0.001,and **** for p<0.0001.

Accelerated GC reactions in CD6 KO cLNs were further confirmed by accelerated transcription of *Aicda*, encoding the AID enzyme which is responsible for B cell somatic hypermutation and antibody isotype switching^34^ (Figure 3C). Therefore, accelerated and enhanced CD4 T_FH_ cell differentiation was followed by increased B cell activation and GC responses. We further confirmed that GC structures were properly forming within CD6 KO cLNs at day 14PI, a time when GCs were easily discernable in WT cLNs, by examining the accumulation of GL7^+^ B cells within the cLN B cell follicle^20, 35, 36^ (Figure 3D). Consistent with accelerated GC somatic hypermutation, mCoV-specific IgG antibodies with increased affinity for mCoV were selectively detected in CD6 KO sera at day 14PI (Figure 3E). Importantly, prior to GC formation we found no difference in the affinity or concentrations of serum mCoV-specific IgG antibodies (Figure 3E-F).

Extended analysis of the GC responses revealed that cLN cellularity was undergoing contraction by day 21 PI. However, contraction of the CD45^+^ cellular population was significantly greater in the CD6 KO cLNs (Supplemental Figure 3A).

Mirroring the CD45^+^ population, contraction of the CD4 T cell population was also greater in CD6 KO mice at day 21PI (Supplemental Figure 3B). However, the proportion of CD4 T_FH_ cells from the total CD4 T cell population was comparable between CD6 KO and WT cLNs (Supplemental Figure 3C). B cell contraction was also trended as increased in the CD6 KO cLNs at day 21PI, but did not reach statistical significance (Supplemental Figure 3D). The relative proportion of GL7^+^ GC B cells within the IgD^-^ B cell population remained comparable between WT and CD6 KO cLNs (Supplemental Figure 3E).

Analysis of the humoral response revealed that high-affinity serum mCoV-specific IgG antibodies were detectable in WT mice by day 21 PI, but affinity was still higher for mCoV-specific IgG antibodies in CD6 KO sera (Supplemental Figure 3F). Intriguingly though, by day 28PI we could no longer detect significant differences in the affinity of serum mCoV-specific IgG antibodies between WT and CD6 KO mice (Supplemental Figure 3F). Semi-quantification of serum mCoV-specific IgG antibodies also revealed that titers were transiently higher in the circulation of CD6 KO mice at day 21 but not at day 28 PI (Supplemental Figure 3F). Whether the enhanced contraction of CD45^+^ cells in CD6 KO cLNs is a direct result of an essential role for CD6 in sustaining antiviral adaptive immune responses, or an indirect of some altered antiviral pathogenesis in CD6 KO mice remains under investigation. Taken together these data demonstrate that the increased CD4 T_FH_ cell differentiation in CD6 KO cLNs was capable of driving accelerated B cell activation and functional GC responses leading to accelerated secretion of high-affinity class-switched virus-specific antibodies.

### Expression of the CD6 intracellular binding protein UBASH3a is suppressed in CD6 KO mice and coincides with stronger TCR signaling

To date, CD6 has predominantly function as a TCR co-stimulatory receptor during T cell activation *in vitro*^6, 7, 37^. Therefore, characterization of the CD6 signaling pathway has primarily identified positive regulators of TCR signaling^6, 7, 37^. However, CD6 has also been shown to directly associate with the negative regulator of T cell activation UBASH3a^6, 7^. Unexpectedly, *Ubash3a* transcription was substantially upregulated in the cLNs of WT mice but was largely abrogated in CD6 KO cLNs in response to mCoV infection (Figure 4A).

**Figure 4:**
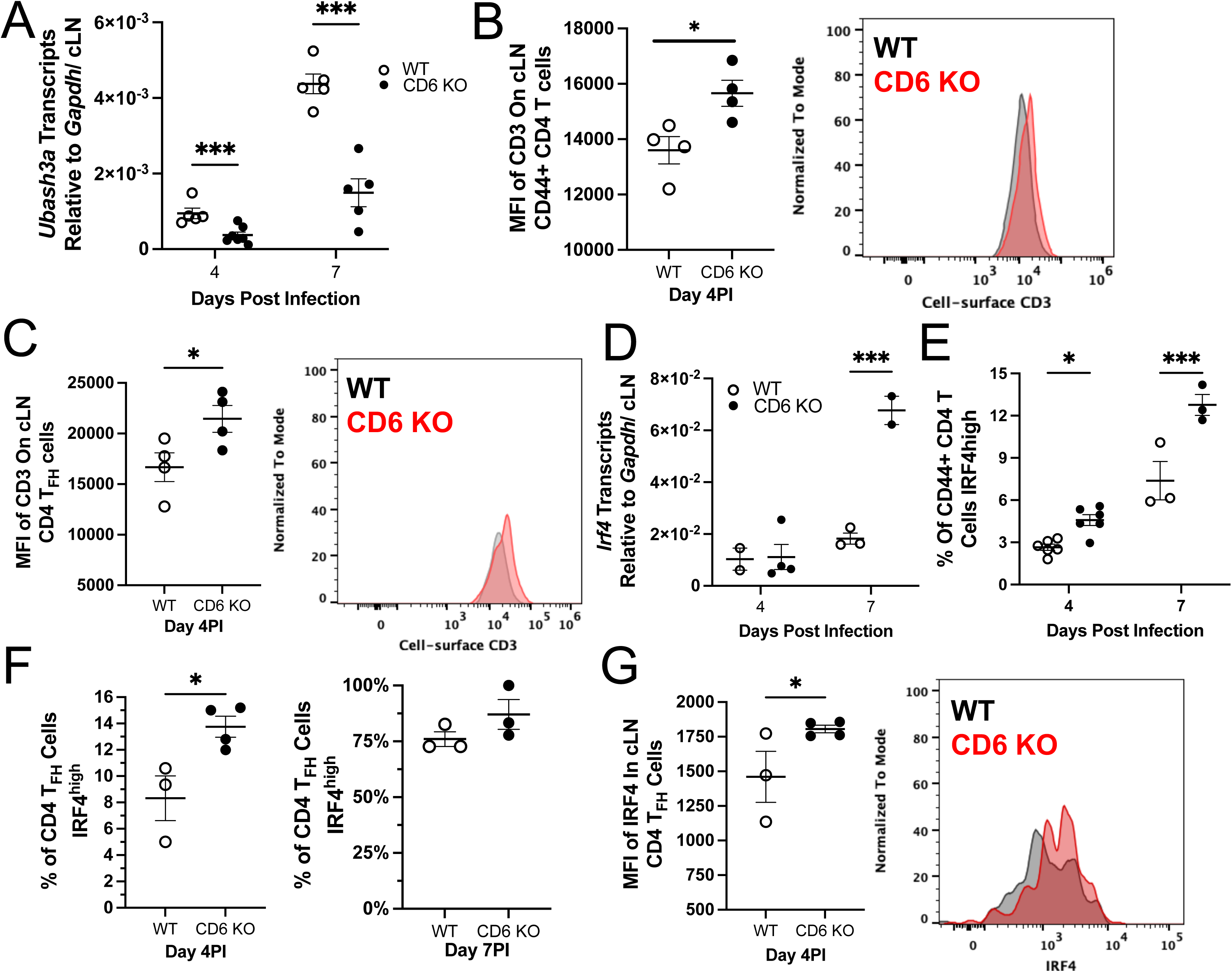
CD6-mediated control of *Ubahs3a* expression corresponds with decreased TCR signal strength. cLNs from mCoV infected WT and CD6 KO mice were isolated at the indicated time points. **A)** *Ubash3a* expression was quantified by qRT-PCR. The MFI of CD3 on **B)** CD44^+^ CD4 T cells and **C)** CD4 T_FH_ cells at day 4PI was used to monitor cell-surface CD3/TCR by flow cytometry. **D)** *Irf4* transcription in the cLNs was quantified by qRT-PCR. The percent of **E)** CD44+ CD4 T cells and **F)** CD4 T_FH_ cells that are IRF4^high^ in the cLN as well as the **G)** MFI of IRF4 on CD44+ CD4 T_FH_ cells was analyzed by flow cytometry. Each data point represents a single mouse and representative histograms are of one WT mouse in grey and one CD6 KO mouse in red. **B, C,** and **H** are data representative of a minimum of 2 experiments. Significance was determined by Two-way ANOVA using a Bonferroni’s post-hoc test (A, D, E) or by unpaired T-Test (B-C and F-G) and denoted as * for p<0.05, ** for p<0.01, *** for p<0.001,and **** for p<0.0001.

We therefore sought to substantiate a functional UBASH3a deficiency in CD6 KO CD4 T cells *in vivo.* While the molecular functions of UBASH3a are poorly delineated, it is established to have weak phosphatase activity^38, 39^ and to negatively regulate cell-surface TCR/CD3 complexes on CD4 T cells^40, 41^. In the absence of a defined mCoV-epitope on the MHCII H-2^q^ background, and thus antigen-specific T cell tetramers, we assessed CD3 expression on the entire CD44^+^ CD4 T cell population, as well as early differentiating CD4 T_FH_ cells. In the absence of CD6, cell-surface CD3 was modestly increased in both total CD44^+^ CD4 T cells (Figure 4B) and CD4 T_FH_ cells (Figure 4C).

While the degree of cell-surface CD3 elevation was minor, it was consistent with the degree of change observed in UBASH3a knockdown studies^40^. These results supported that CD6 may utilize UBASH3a to suppress T cell activation in cLNs during mCoV infection.

As UBASH3a is a negative regulator of TCR signal strength, we next assessed differences in the strength of TCR signal in cLN CD4 T cells from infected CD6 KO and WT mice by measuring the transcription factor IRF4, an established dose-dependent readout of the TCR signal strength^1, 4, 42–45^. Given the increase in GC responses *Irf4* transcripts were unsurprisingly highly elevated by day 7 PI in CD6 KO compared to WT cLNs (Figure 4D). Target analysis of CD4 T cell populations by flow cytometry confirmed that a higher percentage of CD6 KO CD44^+^ CD4 T cells express high levels of IRF4, signifying that CD6 KO T cells had stronger TCR signaling after mCoV infection (Figure 4E). During the early stages of CD4 T_FH_ cell differentiation (day 4PI), CD6 KO CD4 T_FH_ cells also displayed elevated IRF4 expression (Figure 4F-G). However, by day 7PI all of the CD4 T_FH_ cells were IRF4 positive, with most being IRF4^high^ (Figure 4F).

Taken together, these data strongly implicate that CD6 suppresses T cell activation during mCoV infection through UBASH3a mediated negative regulation of the TCR signaling.

### CD6 regulates peripheral, but not CNS, humoral immunity during mCoV induced encephalomyelitis

We next examined the adaptive immune responses within the mCoV infected CNS. Consistent with greater activation and expansion in the cLN, CD4 T cell numbers were elevated in the CD6 KO infected brain at day 7PI (Figure 5A). On the other hand, CD8 T cell infiltration of the brain was not significantly altered at day 7PI (Figure 5B).

**Figure 5:**
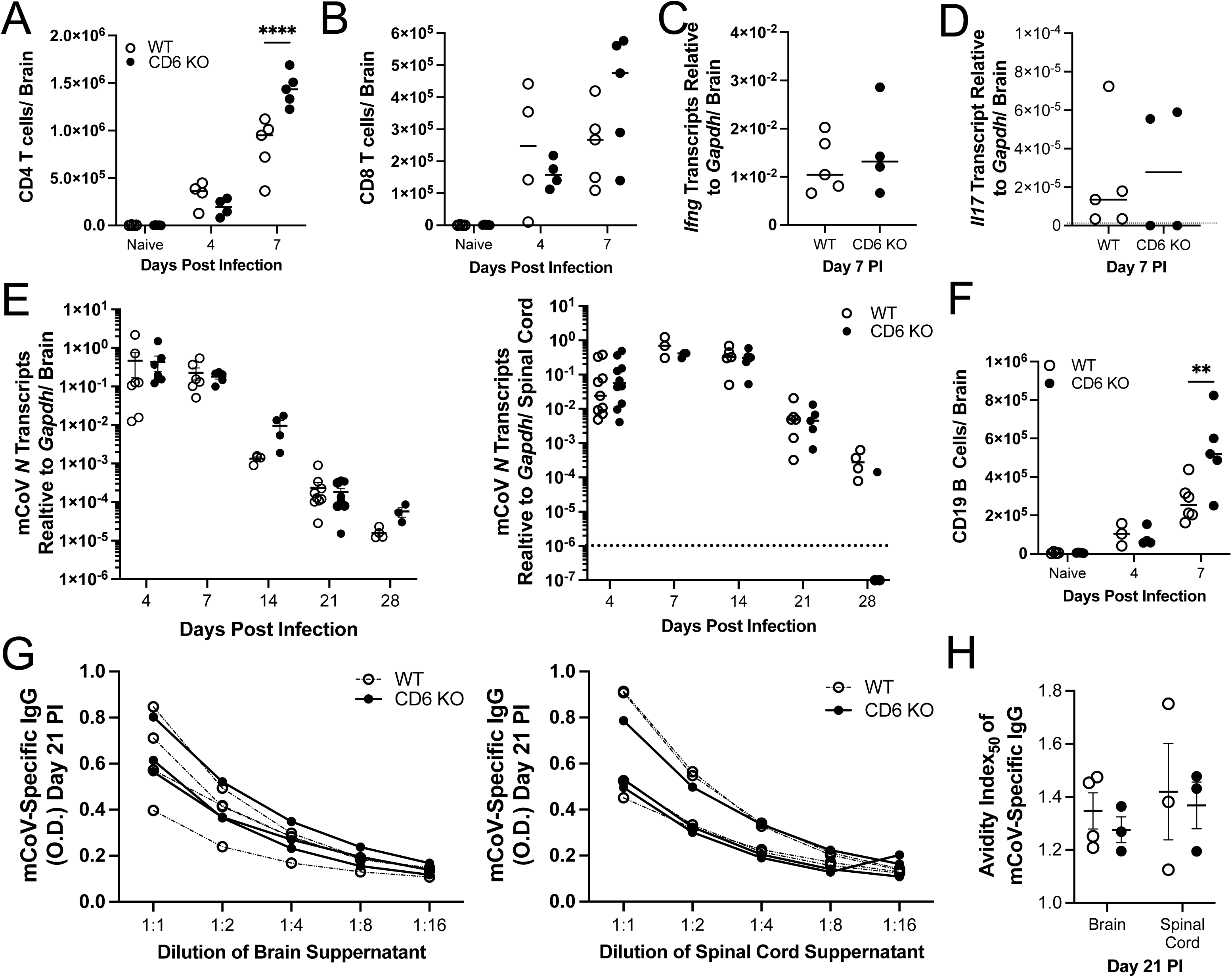
CD6 regulates peripheral, but not CNS, immune responses. CNS tissues were isolated from WT and CD6 KO mice at the indicated time points. The total number of **A)** CD4 T cells and **B)** CD8 T cells were quantified by flow cytometry. mRNA transcripts of **C)** *Ifng,* **D)** *Il17,* and **E)** viral *N* gene were quantified in the indicated CNS tissues. The total number of **F)** CD19 B cells in the brain was quantified by flow cytometry. mCoV-specific IgG antibodies in the CNS tissues were **G)** semi-quantified and **H)** measured for affinity using an avidity index_50_ assay. Each data point represents a single mouse. Significance was determined by A-B) Two-way ANOVA using a Bonferroni’s post-hoc test, C-D) unpaired T-Test and E-G) Two-way ANOVA using a Bonferroni’s post-hoc test and denoted as * for p<0.05, ** for p<0.01, *** for p<0.001,and **** for p<0.0001.

*Ifng* transcripts were similar between WT and CD6 KO brains (Figure 5C) and *Il17* transcripts remained undetectable (Figure 5D). The elevated total number of CD4 T cells in CD6 KO infected brains was thus not associated with overt differences in T cell effector activity. Furthermore, comparable antiviral effector T cell responses were supported by congruent kinetics of viral control between WT and CD6 KO brains and spinal cords as measured by viral nucleocapsid (N) specific transcripts (Figure 5E).

The total number of B cells in the CD6 KO mCoV infected brains was also elevated at day 7PI (Figure 5F). This was notable as most of the B cells in the brain at this early timepoint are IgD^+^IgM^+^ B cells that migrate to the CNS in response to the inflammation^20–21^. More surprisingly, analysis of supernatants from dissociated WT and CD6 KO brain and spinal cord tissue revealed similar levels of virus-specific IgG antibodies across multiple dilutions as well as comparable mCoV affinity at day 21PI (Figure 5G-H). Therefore, while CD6 KO mice transiently had greater mCoV-specific IgG responses in the periphery at day 21PI (Supplemental Figure 3 F), there were no detectable changes within the infected CNS at this time point. Thus, these data implicated that CD6 predominantly regulates the peripheral CD4 T cell and humoral responses at the priming site, with minimal impact in the infected CNS during mCoV infection.

## Discussion

The dual functions of CD6 as a positive and negative regulator of T cell activation are well established, but its influence on T cell responses during viral infections have not been studied. In this report, we investigated how the absence of CD6 affects T cell activation using a neurotropic mCoV model, in which CD4 and CD8 T cells are both essential to control infectious virus. Our results demonstrate that CD6 acts as a negative regulator of CD4 T cell activation in CNS draining cLN following virus infection. In addition, we have discovered a previously unrecognized role for CD6 in limiting CD4T_FH_ cell differentiation and, by extension, delaying GC responses. These novel findings have significant clinical implications for patients receiving therapeutic CD6 blocking antibody treatments.

The function of CD6 as a negative regulator of T cell activation during mCoV encephalomyelitis stands in stark contrast to its co-stimulatory role in multiple autoimmune models including autoimmune encephalitis and autoimmune uveitis ^5–6, 15–16, 46–47^. Importantly, the opposing functions of CD6 in CD4 T cell activation are not easily attributed to murine intrinsic factors, as the CD6 KO mice used herein were generated from the same colony and housed in the same facility as in the above mentioned autoimmune studies^15–16^. Unfortunately, further comparison between autoimmune and virus models is complicated by numerous factors; In the autoimmune encephalomyelitis and uveitis models T cell activation is dependent on immunization with self-peptide or antigen in adjuvant resulting in the induction of both Th1 and Th17 CD4 T cells^15, 46^, whereas virus-specific T cells are activated by replicating virus and presentation of viral antigen generating an exclusive Th1 response^17–20^. These models also utilize distinct innate immune scavenging, pattern recognition receptors, and antigen presenting cells, all of which also contribute to the outcome of T cell activation^48–50^. It is thus reasonable that such distinct innate input signals *in viv*o would influence differential expression of the extracellular CD6 ligands and/ or CD6 intracellular interacting proteins, leading to distinct outcomes of CD6 signaling.

To this end, analysis of the established CD6 intracellular signaling proteins identified UBASH3A as a primary candidate responsible for CD6 mediated dampening of CD4 T cell activation following mCoV infection. *Ubash3a* mRNA transcripts increased in the cLNs of WT mice, while upregulation was drastically impaired in CD6 KO cLNS. Consistent with UBASH3a knockdown studies and diminished UBASH3a activity, cLN CD4 T cells from CD6 KO mice had modest, but significantly, increased accumulation of cell-surface CD3^40^. While these fairly minor changes in increased cell-surface CD3/TCR complexes are unlikely to result in enhanced T cell activation, as few as 500 TCR complexes have been implicated to be sufficient for T cell activation *in vivo*^51^.

UBASH3a is incompletely characterized, but is known to suppress TCR signaling, at least in part through suppression of ZAP-70 signaling in a ubiquitin and phosphatase-dependent manner^39, 52–53^. Since TCR signaling is established to directly upregulate IRF4 in a dose-dependent manner in both CD8 and CD4 T cells^43, 44, 45^, IRF4 expression was used as a readout to measure how loss of UBASH3a in CD6 KO mice affected the TCR signal strength. Indeed, IRF4 protein levels were higher in CD6 KO CD4T cells compared to WT CD4 T cells from the cLNs, confirming that the absence of CD6 produced a stronger TCR signal. Overall, the novel finding that CD6 influences *Ubash3a* expression in the mCoV infection model implicates that UBASH3a exerts inhibitory effects on TCR signaling through CD6. Although UBASH3a is known to suppress T cell activation and proliferation *in vitro* and in autoimmune diabetes^54^, UBASH3a regulation has not been explored in studies focusing on CD6. Therefore, analysis of UBASH3a in autoimmune diseases where CD6 acts as a positive regulator of T cell activation may shed light on the pathogenic role of CD6.

Unexpectedly, CD6 KO CD4 T cells also showed preferential differentiation into CD4 T_FH_ effectors. While TCR-signal strength is an established determinate of T helper cell differentiation^1–4^, mechanistic studies to assess how CD6 signaling proteins, including UBASH3a, regulate CD4 T_FH_ cell differentiation will require the development of complex *in vitro* assays that mimic T cell activation during mCoV infection *in vivo*.

Nevertheless, the accelerated and enhanced T_FH_ cell development correlated with earlier and elevated titers of high-affinity class-switched virus-specific antibodies in the serum of CD6 KO mice. Of note, the difference in the total number of CD44^+^ CD4 T cells could not be completely explained by the magnitude of change in CD4 T_FH_ cells. As we were unable to detect changes in other CD4 T helper cells, and there are indications that both CD6 KO and UBASH3a KO T cells may have a lower threshold for activation homeostatically, it is likely that there is some degree of non-specific T cell recruitment and activation contributing to the greater accumulation of CD4 T cell in the CD6 KO cLNs^15, 60^.

Distinct from the periphery, we found no difference in mCoV-specific IgG antibodies within the CNS itself. We are currently investigating whether early changes in the CD6 KO cLNs alters the differentiation, survival, or migratory capacity acquired of antibody secreting cells during GC differentiation^55^. To this end, it is worth noting that the predominant ligand of CD6, CD166, was found to be essential for pathogenic B cells to infiltrate the CNS during experimental autoimmune encephalitis^56^. Interestingly though, while CD6 appears to facilitate T cell migration in autoimmune encephalomyelitis^15^, it was redundant for T cell infiltration into the virally infected CNS. This redundancy may also explain why CD6 was essential to sustain dendritic cell-T cell interaction during antigen presentation *in vitro*, but not in this *in vivo* setting^61^.

Overall, these data indicate that the clinically used CD6 blocking antibody treatments may be beneficial to antiviral immunity. A limitation of the mCoV model is that acute viral replication in the CNS is controlled by T cells and not the humoral response, which was reflected in the similar kinetics of virus control between the WT and CD6 KO CNS. Therefore, the biological significance of the accelerated antiviral-humoral response in the absence of CD6 may be more readily revealed in a model where GC-derived humoral responses are essential to prevent viral dissemination to the CNS. CD6 may also potentially be exploited during the administration of traditional and mRNA-based vaccines. As Itolizumab has been in use in India since 2013 and given the number of clinical trials ongoing during the COIVD-19 pandemic, it may be feasible to examine antiviral humoral responses during vaccination as well as primary SARS-CoV2 infections in patients that had been receiving CD6 monoclonal blocking antibody therapies^8, 12, 57–59^.

In summary, we have identified novel roles for CD6 as a negative regulator of both CD4 T cell and GC-derived antiviral humoral responses. CD6 inhibition of these responses appears to be T cell intrinsic and associated with a deficit in UBASH3a mediated suppression of CD4 T cell TCR signal. The number of ongoing clinical trials examining the efficacy of CD6 blockade necessitates further interrogation of its role during viral infections and vaccination, especially given the context dependent role of CD6 in T cell activation.

## Acknowledgements

This work was supported by the Nation Institute of Health R01 NS086299 (C.B.) and R01 EY034077 (F.L)

## Author Contributions

A.C-B. and C.C.B. Study conception and design; A.C-B. Acquired and Analyzed data; A.C-B. wrote the manuscript; C.C.B. and F.L. Edited the manuscript; F.L and C.C.B., Providing funding/ reagents and supervision.

---

**Supplemental Figure 1: Representative gating strategy of mCoV infected cLNs at day 7PI. S**inglet cells that fell into a lymphocyte FSC, SSC gate were analyzed for cell viability using a live/dead dye. CD45+ live cells were then first A) analyzed for CD3e expression followed by CD4 or CD8 expression. CD4 expressing T cells were then analyzed for activation CD44 and T_FH_ cells by CXCR5 and PD1 co-expression. B) CD45+ cells were also analyzed for CD19, followed by GL7 expression or IgD expression. Finally, IgD-B cells were examined for CD138.

**Supplemental Figure 2: CD6 is expressed on CD4 and CD8 T cells.** A) cLNs and B) brains from naïve WT (n=4) and CD6 KO (n=2) mice were analyzed by flow cytometry for CD6 expression on the indicated population. Representative histograms of CD6 expression in the naïve cLNs are shown in the left panel of (A). C) Representative flow cytometry plots from the naïve cLNs (left) and brain (right) demonstrating no detectable CD6 expression on CD45^-^ cells. D) Representative histograms of cLNs were taken at day 4 (left) and 7 (right) PI of CD6 expression on live cells with T cells gated out. E) CD6 expression on activated T cells at day 4PI in the cLNs was confirmed by flow cytometry. All cells were identified by flow cytometery using the gating scheme depicted in Supplemental Figure 1. Each data point represents an individual mouse and representative histograms and flow plots are from an individual WT (red) and CD6 KO (grey) mouse.

**Supplemental Figure 3: Contraction in the cLNs is accelerated in CD6 KO mice.** cLNs from WT and CD6 KO mice were analyzed by flow cytometry at the indicated time point(s) to quantify **A)** total CD45^+^ cells (simplified graph of day 21PI alone on the right to illustrate the difference in contraction), **B)** total CD4 T cells **C)** the fraction of CD4 T cells that are CD4 T_FH_ cells, **D)** total CD19 cells, and **E)** the percent of IgD^-^ B cells that are GC B cells. At days 21 and 28 PI the mCoV-specific IgG antibody **F)** avidity index_50_ (left) and titers (right) were measured across multiple experiments. For all graphs each data points represents an individual mouse. Significance was determined by Two-way ANOVA using a Bonferroni’s post-hoc test or an unpaired T test and denoted as * for p<0.05, ** for p<0.01, *** for p<0.001,and **** for p<0.0001.

## References

1. Tubo NJ, Jenkins MK. TCR signal quantity and quality in CD4^+^ T cell differentiation. Trends Immunol. 2014 Dec;35(12):591–596. doi: 10.1016/j.it.2014.09.008. Epub 2014 Oct 22. PMID: 25457838; PMCID: PMC4406772.

2. Künzli M, Reuther P, Pinschewer DD, King CG. Opposing effects of T cell receptor signal strength on CD4 T cells responding to acute versus chronic viral infection. Elife. 2021 Mar 8;10:e61869. doi: 10.7554/eLife.61869. PMID: 33684030; PMCID: PMC7943189.

3. Snook JP, Kim C, Williams MA. TCR signal strength controls the differentiation of CD4^+^ effector and memory T cells. Sci Immunol. 2018 Jul 20;3(25):eaas9103. doi: 10.1126/sciimmunol.aas9103. PMID: 30030369; PMCID: PMC6126666.

4. Bhattacharyya ND, Feng CG. Regulation of T Helper Cell Fate by TCR Signal Strength. Front Immunol. 2020 May 19;11:624. doi: 10.3389/fimmu.2020.00624. PMID: 32508803; PMCID: PMC7248325.

5. Gonçalves CM, Henriques SN, Santos RF, Carmo AM. CD6, a Rheostat-Type Signalosome That Tunes T Cell Activation. Front Immunol. 2018 Dec 18;9:2994. doi: 10.3389/fimmu.2018.02994. PMID: 30619347; PMCID: PMC6305463.

6. Mori D, Grégoire C, Voisinne G, Celis-Gutierrez J, Aussel R, Girard L, Camus M, Marcellin M, Argenty J, Burlet-Schiltz O, Fiore F, Gonzalez de Peredo A, Malissen M, Roncagalli R, Malissen B. The T cell CD6 receptor operates a multitask signalosome with opposite functions in T cell activation. J Exp Med. 2021 Feb 1;218(2):e20201011. doi: 10.1084/jem.20201011. PMID: 33125054; PMCID: PMC7608068.

7. Voisinne G, Locard-Paulet M, Froment C, Maturin E, Menoita MG, Girard L, Mellado V, Burlet-Schiltz O, Malissen B, Gonzalez de Peredo A, Roncagalli R. Kinetic proofreading through the multi-step activation of the ZAP70 kinase underlies early T cell ligand discrimination. Nat Immunol. 2022 Sep;23(9):1355–1364. doi: 10.1038/s41590-022-01288-x. Epub 2022 Aug 31. PMID: 36045187; PMCID: PMC9477740.

8. Krupashankar DS, Dogra S, Kura M, Saraswat A, Budamakuntla L, Sumathy TK, Shah R, Gopal MG, Narayana Rao T, Srinivas CR, Bhat R, Shetty N, Manmohan G, Sai Krishna K, Padmaja D, Pratap DV, Garg V, Gupta S, Pandey N, Khopkar U, Montero E, Ramakrishnan MS, Nair P, Ganapathi PC. Efficacy and safety of itolizumab, a novel anti-CD6 monoclonal antibody, in patients with moderate to severe chronic plaque psoriasis: results of a double-blind, randomized, placebo-controlled, phase-III study. J Am Acad Dermatol. 2014 Sep;71(3):484–92. doi: 10.1016/j.jaad.2014.01.897. Epub 2014 Apr 2. PMID: 24703722.

9. De Jager PL, Jia X, Wang J, de Bakker PI, Ottoboni L, Aggarwal NT, Piccio L, Raychaudhuri S, Tran D, Aubin C, Briskin R, Romano S; International MS Genetics Consortium; Baranzini SE, McCauley JL, Pericak-Vance MA, Haines JL, Gibson RA, Naeglin Y, Uitdehaag B, Matthews PM, Kappos L, Polman C, McArdle WL, Strachan DP, Evans D, Cross AH, Daly MJ, Compston A, Sawcer SJ, Weiner HL, Hauser SL, Hafler DA, Oksenberg JR. Meta-analysis of genome scans and replication identify CD6, IRF8 and TNFRSF1A as new multiple sclerosis susceptibility loci. Nat Genet. 2009 Jul;41(7):776–82. doi: 10.1038/ng.401. Epub 2009 Jun 14. PMID: 19525953; PMCID: PMC2757648.

10. Zheng M, Zhang L, Yu H, Hu J, Cao Q, Huang G, Huang Y, Yuan G, Kijlstra A, Yang P. Genetic polymorphisms of cell adhesion molecules in Behcet’s disease in a Chinese Han population. Sci Rep. 2016 Apr 25;6:24974. doi: 10.1038/srep24974. PMID: 27108704; PMCID: PMC4842956.

11. Consuegra-Fernández M, Julià M, Martínez-Florensa M, Aranda F, Català C, Armiger-Borràs N, Arias MT, Santiago F, Guilabert A, Esteve A, Muñoz C, Ferrándiz C, Carrascosa JM, Pedrosa E, Romaní J, Alsina M, Mascaró-Galy JM, Lozano F. Genetic and experimental evidence for the involvement of the CD6 lymphocyte receptor in psoriasis. Cell Mol Immunol. 2018 Oct;15(10):898–906. doi: 10.1038/cmi.2017.119. Epub 2017 Dec 11. PMID: 29225340; PMCID: PMC6207571.

12. Rodríguez PC, Prada DM, Moreno E, Aira LE, Molinero C, López AM, Gómez JA, Hernández IM, Martínez JP, Reyes Y, Milera JM, Hernández MV, Torres R, Avila Y, Barrese Y, Viada C, Montero E, Hernández P. The anti-CD6 antibody itolizumab provides clinical benefit without lymphopenia in rheumatoid arthritis patients: results from a 6-month, open-label Phase I clinical trial. Clin Exp Immunol. 2018 Feb;191(2):229–239. doi: 10.1111/cei.13061. Epub 2017 Nov 16. PMID: 28963724; PMCID: PMC5758380.

13. Ruth JH, Gurrea-Rubio M, Athukorala KS, Rasmussen SM, Weber DP, Randon PM, Gedert RJ, Lind ME, Amin MA, Campbell PL, Tsou PS, Mao-Draayer Y, Wu Q, Lanigan TM, Keshamouni VG, Singer NG, Lin F, Fox DA. CD6 is a target for cancer immunotherapy. JCI Insight. 2021 Mar 8;6(5):e145662. doi: 10.1172/jci.insight.145662. PMID: 33497367; PMCID: PMC8021120.

14. Gurrea-Rubio M, Wu Q, Amin MA, Tsou PS, Campbell PL, Amarista CI, Ikari Y, Brodie WD, Mattichak MN, Muraoka S, Randon PM, Lind ME, Ruth JH, Mao-Draayer Y, Ding S, Shen X, Cooney LA, Lin F, Fox DA. Activation of cytotoxic lymphocytes through CD6 enhances killing of cancer cells. Cancer Immunol Immunother. 2024 Jan 27;73(2):34. doi: 10.1007/s00262-023-03578-1. PMID: 38280067; PMCID: PMC10821976.

15. Li Y, Singer NG, Whitbred J, Bowen MA, Fox DA, Lin F. CD6 as a potential target for treating multiple sclerosis. Proc Natl Acad Sci U S A. 2017 Mar 7;114(10):2687–2692. doi: 10.1073/pnas.1615253114. Epub 2017 Feb 16. PMID: 28209777; PMCID: PMC5347585.

16. Henriques, S.N., Oliveira, L., Santos, R.F. et al. CD6-mediated inhibition of T cell activation via modulation of Ras. Cell Commun Signal 20, 184 (2022). 10.1186/s12964-022-00998-x

17. Cowley TJ, Weiss SR. Murine coronavirus neuropathogenesis: determinants of virulence. J Neurovirol. 2010 Nov;16(6):427–34. doi: 10.3109/13550284.2010.529238. Epub 2010 Nov 12. PMID: 21073281; PMCID: PMC3153983.

18. Stohlman SA, Bergmann CC, Lin MT, Cua DJ, Hinton DR (1998). CTL effector function within the central nervous system requires CD4+ T cells. J Immunol 160: 2896–2904.

19. Sussman MA, Shubin RA, Kyuwa S, Stohlman SA (1989). T-cell-mediated clearance of mouse hepatitis virus strain JHM from the central nervous system. J Virol 63: 3051–3061.

20. Cardani-Boulton A, Boylan BT, Stetsenko V, Bergmann CC. B cells going viral in the CNS: Dynamics, complexities, and functions of B cells responding to viral encephalitis. Immunol Rev. 2022 Oct;311(1):75–89. doi: 10.1111/imr.13124. Epub 2022 Aug 19. PMID: 35984298; PMCID: PMC9804320.

21. Atkinson JR, Bergmann CC. Protective Humoral Immunity in the Central Nervous System Requires Peripheral CD19-Dependent Germinal Center Formation following Coronavirus Encephalomyelitis. J Virol. 2017 Nov 14;91(23):e01352–17. doi: 10.1128/JVI.01352-17. PMID: 28931676; PMCID: PMC5686739.

22. Lin MT, Hinton DR, Marten NW, Bergmann CC, Stohlman SA (1999). Antibody prevents virus reactivation within the central nervous system. J Immunol 162:7358–7368.

23. Matthews AE, Weiss SR, Shlomchik MJ, Hannum LG, Gombold JL, Paterson Y. Antibody is required for clearance of infectious murine hepatitis virus A59 from the central nervous system, but not the liver. J Immunol. 2001 Nov 1;167(9):5254–63. doi: 10.4049/jimmunol.167.9.5254. PMID: 11673540.

24. Das Sarma J, Scheen E, Seo SH, Koval M, Weiss SR. Enhanced green fluorescent protein expression may be used to monitor murine coronavirus spread in vitro and in the mouse central nervous system. J Neurovirol. 2002 Oct;8(5):381–91. doi: 10.1080/13550280260422686. PMID: 12402164; PMCID: PMC7095158.

25. Akache B, Stark FC, McCluskie MJ. Measurement of Antigen-Specific IgG Titers by Direct ELISA. Methods Mol Biol. 2021;2183:537–547. doi: 10.1007/978-1-0716-0795-4_31. PMID: 32959266.

26. Amber Cardani-Boulton, Sun-Sang J. Sung, William A. Petri, Young S. Hahn, Thomas J. Braciale; Leptin Receptor Deficiency Impairs Lymph Node Development and Adaptive Immune Response. J Immunol 2024; ji2100985. 10.4049/jimmunol.2100985

27. Metcalf TU, Griffin DE. Alphavirus-induced encephalomyelitis: antibody-secreting cells and viral clearance from the nervous system. J Virol. 2011 Nov;85(21):11490–501. doi: 10.1128/JVI.05379-11. Epub 2011 Aug 24. PMID: 21865385; PMCID: PMC3194963.

28. Pullen GR, Fitzgerald MG, Hosking CS. Antibody avidity determination by ELISA using thiocyanate elution. J Immunol Methods. 1986 Jan 22;86(1):83–7. doi: 10.1016/0022-1759(86)90268-1. PMID: 3944471.

29. Macdonald RA, Hosking CS, Jones CL. The measurement of relative antibody affinity by ELISA using thiocyanate elution. J Immunol Methods. 1988 Feb 10;106(2):191–4. doi: 10.1016/0022-1759(88)90196-2. PMID: 3339255.

30. Enyindah-Asonye G, Li Y, Xin W, Singer NG, Gupta N, Fung J, Lin F. CD6 Receptor Regulates Intestinal Ischemia/Reperfusion-induced Injury by Modulating Natural IgM-producing B1a Cell Self-renewal. J Biol Chem. 2017 Jan 13;292(2):661–671. doi: 10.1074/jbc.M116.749804. Epub 2016 Dec 1. PMID: 27909060; PMCID: PMC5241740.

31. Braun M, Müller B, ter Meer D, Raffegerst S, Simm B, Wilde S, Spranger S, Ellwart J, Mosetter B, Umansky L, Lerchl T, Schendel DJ, Falk CS. The CD6 scavenger receptor is differentially expressed on a CD56 natural killer cell subpopulation and contributes to natural killer-derived cytokine and chemokine secretion. J Innate Immun. 2011;3(4):420–34. doi: 10.1159/000322720. Epub 2010 Dec 18. PMID: 21178331.

32. Choi J, Crotty S. Bcl6-Mediated Transcriptional Regulation of Follicular Helper T cells (TFH). Trends Immunol. 2021 Apr;42(4):336–349. doi: 10.1016/j.it.2021.02.002. Epub 2021 Mar 1. PMID: 33663954; PMCID: PMC8021443.

33. Szabo SJ, Kim ST, Costa GL, Zhang X, Fathman CG, Glimcher LH. A novel transcription factor, T-bet, directs Th1 lineage commitment. Cell. 2000 Mar 17;100(6):655–69. doi: 10.1016/s0092-8674(00)80702-3. PMID: 10761931.

34. Okazaki IM, Kinoshita K, Muramatsu M, Yoshikawa K, Honjo T. The AID enzyme induces class switch recombination in fibroblasts. Nature. 2002 Mar 21;416(6878):340-5. doi: 10.1038/nature727. Epub 2002 Mar 3. PMID: 11875397.

35. Cyster JG, Allen CDC. B Cell Responses: Cell Interaction Dynamics and Decisions. Cell. 2019 Apr 18;177(3):524–540. doi: 10.1016/j.cell.2019.03.016. PMID: 31002794; PMCID: PMC6538279.

36. Laidlaw BJ, Cyster JG. Transcriptional regulation of memory B cell differentiation. Nat Rev Immunol. 2021 Apr;21(4):209–220. doi: 10.1038/s41577-020-00446-2. Epub 2020 Oct 6. PMID: 33024284; PMCID: PMC7538181.

37. Roncagalli R, Hauri S, Fiore F, Liang Y, Chen Z, Sansoni A, Kanduri K, Joly R, Malzac A, Lähdesmäki H, Lahesmaa R, Yamasaki S, Saito T, Malissen M, Aebersold R, Gstaiger M, Malissen B. Quantitative proteomics analysis of signalosome dynamics in primary T cells identifies the surface receptor CD6 as a Lat adaptor-independent TCR signaling hub. Nat Immunol. 2014 Apr;15(4):384–392. doi: 10.1038/ni.2843. Epub 2014 Mar 2. PMID: 24584089; PMCID: PMC4037560.

38. Mikhailik A, Ford B, Keller J, Chen Y, Nassar N, Carpino N. A phosphatase activity of Sts-1 contributes to the suppression of TCR signaling. Mol Cell. 2007 Aug 3;27(3):486–97. doi: 10.1016/j.molcel.2007.06.015. PMID: 17679096; PMCID: PMC2709417.

39. Luis BS, Carpino N. Insights into the suppressor of T-cell receptor (TCR) signaling-1 (Sts-1)-mediated regulation of TCR signaling through the use of novel substrate-trapping Sts-1 phosphatase variants. FEBS J. 2014 Feb;281(3):696–707. doi: 10.1111/febs.12615. Epub 2013 Dec 12. PMID: 24256567; PMCID: PMC3968691.

40. Ge Y, Paisie TK, Chen S, Concannon P. UBASH3A Regulates the Synthesis and Dynamics of TCR-CD3 Complexes. J Immunol. 2019 Dec 1;203(11):2827–2836. doi: 10.4049/jimmunol.1801338. Epub 2019 Oct 28. PMID: 31659016; PMCID: PMC6938261.

41. Voisinne G, García-Blesa A, Chaoui K, Fiore F, Bergot E, Girard L, Malissen M, Burlet-Schiltz O, Gonzalez de Peredo A, Malissen B, Roncagalli R. Co-recruitment analysis of the CBL and CBLB signalosomes in primary T cells identifies CD5 as a key regulator of TCR-induced ubiquitylation. Mol Syst Biol. 2016 Jul 29;12(7):876. doi: 10.15252/msb.20166837. PMID: 27474268; PMCID: PMC4965873.

42. Yao S, Buzo BF, Pham D, Jiang L, Taparowsky EJ, Kaplan MH, Sun J. Interferon regulatory factor 4 sustains CD8(+) T cell expansion and effector differentiation. Immunity. 2013 Nov 14;39(5):833–45. doi: 10.1016/j.immuni.2013.10.007. Epub 2013 Nov 7. PMID: 24211184; PMCID: PMC3855863.

43. Wu H, Witzl A, Ueno H. Assessment of TCR signal strength of antigen-specific memory CD8^+^ T cells in human blood. Blood Adv. 2019 Jul 23;3(14):2153–2163. doi: 10.1182/bloodadvances.2019000292. PMID: 31320320; PMCID: PMC6650739.

44. Allison KA, Sajti E, Collier JG, Gosselin D, Troutman TD, Stone EL, Hedrick SM, Glass CK. Affinity and dose of TCR engagement yield proportional enhancer and gene activity in CD4+ T cells. Elife. 2016 Jul 4;5:e10134. doi: 10.7554/eLife.10134. PMID: 27376549; PMCID: PMC4931909.

45. Man K, Miasari M, Shi W, Xin A, Henstridge DC, Preston S, Pellegrini M, Belz GT, Smyth GK, Febbraio MA, Nutt SL, Kallies A. The transcription factor IRF4 is essential for TCR affinity-mediated metabolic programming and clonal expansion of T cells. Nat Immunol. 2013 Nov;14(11):1155–65. doi: 10.1038/ni.2710. Epub 2013 Sep 22. Erratum in: Nat Immunol. 2014 Sep;15(9):894. PMID: 24056747.

46. Zhang L, Li Y, Qiu W, Bell BA, Dvorina N, Baldwin WM 3rd, Singer N, Kern T, Caspi RR, Fox DA, Lin F. Targeting CD6 for the treatment of experimental autoimmune uveitis. J Autoimmun. 2018 Jun;90:84–93. doi: 10.1016/j.jaut.2018.02.004. Epub 2018 Feb 19. PMID: 29472120; PMCID: PMC5949263

47. Li Y, Ruth JH, Rasmussen SM, Athukorala KS, Weber DP, Amin MA, Campbell PL, Singer NG, Fox DA, Lin F. Attenuation of Murine Collagen-Induced Arthritis by Targeting CD6. Arthritis Rheumatol. 2020 Sep;72(9):1505–1513. doi: 10.1002/art.41288. Epub 2020 Aug 14. PMID: 32307907; PMCID: PMC7745675.

48. Marta M, Meier UC, Lobell A. Regulation of autoimmune encephalomyelitis by toll-like receptors. Autoimmun Rev. 2009 May;8(6):506–9. doi: 10.1016/j.autrev.2009.01.006. Epub 2009 Jan 27. PMID: 19211042.

49. Zalinger ZB, Elliott R, Rose KM, Weiss SR. MDA5 Is Critical to Host Defense during Infection with Murine Coronavirus. J Virol. 2015 Dec;89(24):12330–40. doi: 10.1128/JVI.01470-15. Epub 2015 Sep 30. PMID: 26423942; PMCID: PMC4665247.

50. Diebold M, Fehrenbacher L, Frosch M, Prinz M. How myeloid cells shape experimental autoimmune encephalomyelitis: At the crossroads of outside-in immunity. Eur J Immunol. 2023 Oct;53(10):e2250234. doi: 10.1002/eji.202250234. Epub 2023 Aug 21. PMID: 37505465.

51. Labrecque N, Whitfield LS, Obst R, Waltzinger C, Benoist C, Mathis D. How much TCR does a T cell need? Immunity. 2001 Jul;15(1):71–82. doi: 10.1016/s1074-7613(01)00170-4. PMID: 11485739.

52. Yang M, Chen T, Li X, Yu Z, Tang S, Wang C, Gu Y, Liu Y, Xu S, Li W, Zhang X, Wang J, Cao X. K33-linked polyubiquitination of Zap70 by Nrdp1 controls CD8(+) T cell activation. Nat Immunol. 2015 Dec;16(12):1253–62. doi: 10.1038/ni.3258. Epub 2015 Sep 21. Erratum in: Nat Immunol. 2020 Mar;21(3):355. PMID: 26390156.

53. Carpino N, Chen Y, Nassar N, Oh HW. The Sts proteins target tyrosine phosphorylated, ubiquitinated proteins within TCR signaling pathways. Mol Immunol. 2009 Oct;46(16):3224–31. doi: 10.1016/j.molimm.2009.08.015. Epub 2009 Sep 5. PMID: 19733910; PMCID: PMC2757469.

54. Chen YG, Ciecko AE, Khaja S, Grzybowski M, Geurts AM, Lieberman SM. UBASH3A deficiency accelerates type 1 diabetes development and enhances salivary gland inflammation in NOD mice. Sci Rep. 2020 Jul 21;10(1):12019. doi: 10.1038/s41598-020-68956-6. PMID: 32694640; PMCID: PMC7374577.

55. Marques CP, Kapil P, Hinton DR, Hindinger C, Nutt SL, Ransohoff RM, Phares TW, Stohlman SA, Bergmann CC. CXCR3-dependent plasma blast migration to the central nervous system during viral encephalomyelitis. J Virol. 2011 Jul;85(13):6136–47. doi: 10.1128/JVI.00202-11. Epub 2011 Apr 20. PMID: 21507985; PMCID: PMC3126522.

56. Michel L, Grasmuck C, Charabati M, Lécuyer MA, Zandee S, Dhaeze T, Alvarez JI, Li R, Larouche S, Bourbonnière L, Moumdjian R, Bouthillier A, Lahav B, Duquette P, Bar-Or A, Gommerman JL, Peelen E, Prat A. Activated leukocyte cell adhesion molecule regulates B lymphocyte migration across central nervous system barriers. Sci Transl Med. 2019 Nov 13;11(518):eaaw0475. doi: 10.1126/scitranslmed.aaw0475. PMID: 31723036.

57. Caballero A, Filgueira LM, Betancourt J, Sánchez N, Hidalgo C, Ramírez A, Martinez A, Despaigne RE, Escalona A, Diaz H, Meriño E, Ortega LM, Castillo U, Ramos M, Saavedra D, García Y, Lorenzo G, Cepeda M, Arencibia M, Cabrera L, Domecq M, Estévez D, Valenzuela C, Lorenzo P, Sánchez L, Mazorra Z, León K, Crombet T. Treatment of COVID-19 patients with the anti-CD6 antibody itolizumab. Clin Transl Immunology. 2020 Nov 25;9(11):e1218. doi: 10.1002/cti2.1218. PMID: 33304584; PMCID: PMC7688906.

58. Díaz Y, Ramos-Suzarte M, Martín Y, Calderón NA, Santiago W, Viñet O, La O Y, Oyarzábal JPA, Pérez Y, Lorenzo G, Cepeda M, Saavedra D, Mazorra Z, Estevez D, Lorenzo-Luaces P, Valenzuela C, Caballero A, Leon K, Crombet T, Hidalgo CJ. Use of a Humanized Anti-CD6 Monoclonal Antibody (Itolizumab) in Elderly Patients with Moderate COVID-19. Gerontology. 2020;66(6):553–561. doi: 10.1159/000512210. Epub 2020 Oct 26. PMID: 33105142; PMCID: PMC7649683

59. Saavedra D, Añé-Kourí AL, Sánchez N, Filgueira LM, Betancourt J, Herrera C, Manso L, Chávez E, Caballero A, Hidalgo C, Lorenzo G, Cepeda M, Valenzuela C, Ramos M, León K, Mazorra Z, Crombet T. An anti-CD6 monoclonal antibody (itolizumab) reduces circulating IL-6 in severe COVID-19 elderly patients. Immun Ageing. 2020 Nov 14;17(1):34. doi: 10.1186/s12979-020-00207-8. PMID: 33292350; PMCID: PMC7666403.

60. Castro-Sánchez P, Aguilar-Sopeña O, Alegre-Gómez S, Ramirez-Munoz R, Roda-Navarro P. Regulation of CD4^+^ T Cell Signaling and Immunological Synapse by Protein Tyrosine Phosphatases: Molecular Mechanisms in Autoimmunity. Front Immunol. 2019 Jun 26;10:1447. doi: 10.3389/fimmu.2019.01447. PMID: 31297117; PMCID: PMC6607956.

61. Zimmerman AW, Joosten B, Torensma R, Parnes JR, van Leeuwen FN, Figdor CG. Long-term engagement of CD6 and ALCAM is essential for T-cell proliferation induced by dendritic cells. Blood. 2006 Apr 15;107(8):3212–20. doi: 10.1182/blood-2005-09-3881. Epub 2005 Dec 13. PMID: 16352806.

